# Brain-wide functional connectivity artifactually inflates throughout fMRI scans: a problem and solution

**DOI:** 10.1101/2023.09.08.556939

**Authors:** Cole Korponay, Amy C. Janes, Blaise B. Frederick

## Abstract

The fMRI blood oxygen level-dependent (BOLD) signal is a mainstay of neuroimaging assessment of neuronal activity and functional connectivity *in vivo*. Thus, a chief priority is maximizing this signal’s reliability and validity. To this end, the fMRI community has invested considerable effort into optimizing both experimental designs and physiological denoising procedures to improve the accuracy, across-scan reproducibility, and subject discriminability of BOLD-derived metrics like functional connectivity. Despite these advances, we discover that a substantial and ubiquitous defect remains in fMRI datasets: functional connectivity throughout the brain artifactually inflates during the course of fMRI scans – by an average of more than 70% in 15 minutes of scan time - at spatially heterogeneous rates, producing both spatial and temporal distortion of brain connectivity maps. We provide evidence that this inflation is driven by a previously unrecognized time-dependent increase of non-neuronal, systemic low-frequency oscillation (sLFO) blood flow signal during fMRI scanning. This signal is not removed by standard denoising procedures such as independent component analysis (ICA). However, we demonstrate that a specialized sLFO denoising procedure - Regressor Interpolation at Progressive Time Delays (RIPTiDe) - can be added to standard denoising pipelines to significantly attenuate functional connectivity inflation. We confirm the presence of sLFO-driven functional connectivity inflation in multiple independent fMRI datasets – including the Human Connectome Project – as well as across resting-state, task, and sleep-state conditions, and demonstrate its potential to produce false positive findings. Collectively, we present evidence for a previously unknown physiological phenomenon that spatiotemporally distorts estimates of brain connectivity in human fMRI datasets, and present a solution for mitigating this artifact.

## Main

A central question in neuroscience is how the brain’s activity and connections reconfigure as events unfold over time to support adaptive functioning. Such temporally dynamic changes in the neural landscape occur over both developmental (i.e., weeks and years) and immediate (i.e., seconds and minutes)^1^ timescales, and are measurable in humans via functional magnetic resonance imaging (fMRI). The leading fMRI metric for modeling brain connections and how they change over time is functional connectivity (FC). FC quantifies the degree to which the neural activity in different brain regions is coordinated with each other, reflecting the magnitude of their connectedness.

However, the ability to obtain accurate measures of FC and how it changes over time depends on the reliability and validity of the fMRI blood oxygen level-dependent (BOLD) signal - the fundamental input to quantifications of neural activity. There are two major factors that reduce the reliability and validity of neuronal signal estimation with BOLD. The first is that the BOLD signal is a non-specific marker of cerebral hemodynamic activity - it reflects changes in blood flow, volume, and oxygenation related not only to neural activity, but also to cardiac, respiratory, and other physiological processes^2-4^; it is also highly sensitive to head motion^5,6^. To address this challenge, the field has developed a set of computational preprocessing and denoising steps that aim to remove non-neuronal sources of BOLD variability and distill a “cleaned”, neuronally-driven BOLD signal^3,5,7-10^. The second challenge is that even this “cleaned” BOLD signal is inherently noisy^11,12^. To address this challenge, the field has begun to invest heavily in large sample size^11^ and dense sampling^13,14^ approaches that include longer scan durations and more scans per subject to increase signal-to-noise ratios.

In evaluating the reliability of BOLD-based metrics (e.g., FC) and the impact of the aforementioned approaches for improving reliability, the field’s focus has primarily been on between-scan reproducibility and discriminability of subject-specific features^15,16^. This has been measured principally via the intraclass correlation coefficient (ICC)^15,16^. Prior work has shown that increasing the amount of imaging data (via increased scan durations, more numerous scans per subjects, and/or increased sample sizes) helps to increase ICC-indexed between-scan reproducibility and subject discriminability^17,18^.

Considerably less attention has been paid to the “within-scan” reliability of the BOLD signal (i.e., its temporal stability, or “moment-to-moment” reproducibility), which is of particular importance for investigations of neural temporal dynamics. The relative inattention to within-scan reliability may be due to a tacit assumption in the field that in denoised fMRI data, remaining BOLD signal noise is temporally “stationary”, arising from random processes whose characteristics are stable across a scanning session. As a corollary, the neuronal signal is assumed to be the dominant source of spatiotemporally organized, coherent signal variance contributing to measured functional connectivity within scanning acquisitions. However, certain kinds of coherent signal variance would be unlikely to be of neuronal origin. Neuronally-driven structured variance in the BOLD signal tends to be both temporally brief and spatially localized (e.g., intermittent fluctuations in FC strengths and patterns in task-relevant networks in response to task demands). We would not expect neural activity to produce temporally sustained, spatially global BOLD signal trends (e.g., continuously increasing or decreasing brain-wide FC throughout a scan). The presence of such non-neuronally driven temporal distortions in the BOLD signal would confound any analysis attempting to understand how the neural landscape changes over time.

Indeed, we discovered just such an artifact while attempting to examine how repeated exposure to drug-related cues alters FC over time in drug-dependent individuals in a study we performed at McLean Hospital^19^. Findings initially indicated significant FC increases within and across runs in task-related circuits, suggesting an impact of the drug cues. However, we subsequently found that these time-dependent FC increases were 1) present in non-task-related circuits throughout the entire brain, and 2) present throughout the entire brain even during rest. Moreover, we then found that the same spatial pattern of FC inflation was present in multiple independent datasets of non-substance-using individuals, including in the resting-state scans of the Human Connectome Project^20,21^ and in sleep-state scans^9^. Upon further investigating the mechanism driving these changes, we found that the spatiotemporal features of a non-neuronal, physiological source of BOLD variance paralleled those of the FC increases. Finally, we found that removal of this non-neuronal signal via a targeted computational denoising procedure also removed the FC inflation. Together, we present evidence for a physiological phenomenon that occurs across human fMRI datasets that spatiotemporally distorts estimations of brain connectivity, and we also present a solution for mitigating this artifact.

## Results

fMRI data for our n=63 discovery sample of nicotine-dependent individuals (24 female; age 28.7±6.7) was acquired on a Siemens 3T Prisma scanner with a 64-channel head coil at the McLean Imaging Center (MIC)^19^. The scanning session for the MIC sample included a 6-minute resting-state acquisition followed by 5, 5-minute runs of a cue reactivity task. For our primary independent replication sample, we used fMRI data from an n=462 cohort^7^ of non-drug-using individuals (281 female; age 28.66 ±3.65) from the Human Connectome Project (HCP)^20,21^, which was acquired on a Siemens 3T Skyra with a 32-channel head coil. This data included 2 sessions of 2, 14.4-minute resting-state acquisitions. As a secondary replication sample, we used 10-minute resting-state and 15-minute sleep-state scans collected from n=33 healthy subjects (16 female; age 22.1±3.2 years) on a 3T Prisma Siemens Fit scanner with a 20-channel head coil at the Pennsylvania State University (PSU)^9^.

Main analyses for all samples used fully preprocessed and independent component analysis (ICA)-denoised resting-state fMRI data. ICA-based denoising for the MIC and PSU resting-state and sleep-state data was performed via FSL’s ICA-AROMA^22^. ICA-FIX^8^ data - among the most widely used analysis-ready resting-state data in the field - was used for the HCP sample.

### Discovery Sample: Brain-Wide Functional Connectivity Inflation During fMRI Scans

Initially, we sought to investigate how functional connectivity between the insula and striatum - two regions strongly implicated in nicotine dependence^19,23^ - changed over time during the course of a task that presents nicotine-related images and is known to induce craving^19^. We observed (**Fig. 1a**) that all insula subregions^24^ displayed significant (*p*<0.05) run-over-run FC strength increases with all striatal subregions (defined by the Yeo 7-network parcellation^25,26^).

**Figure 1.**
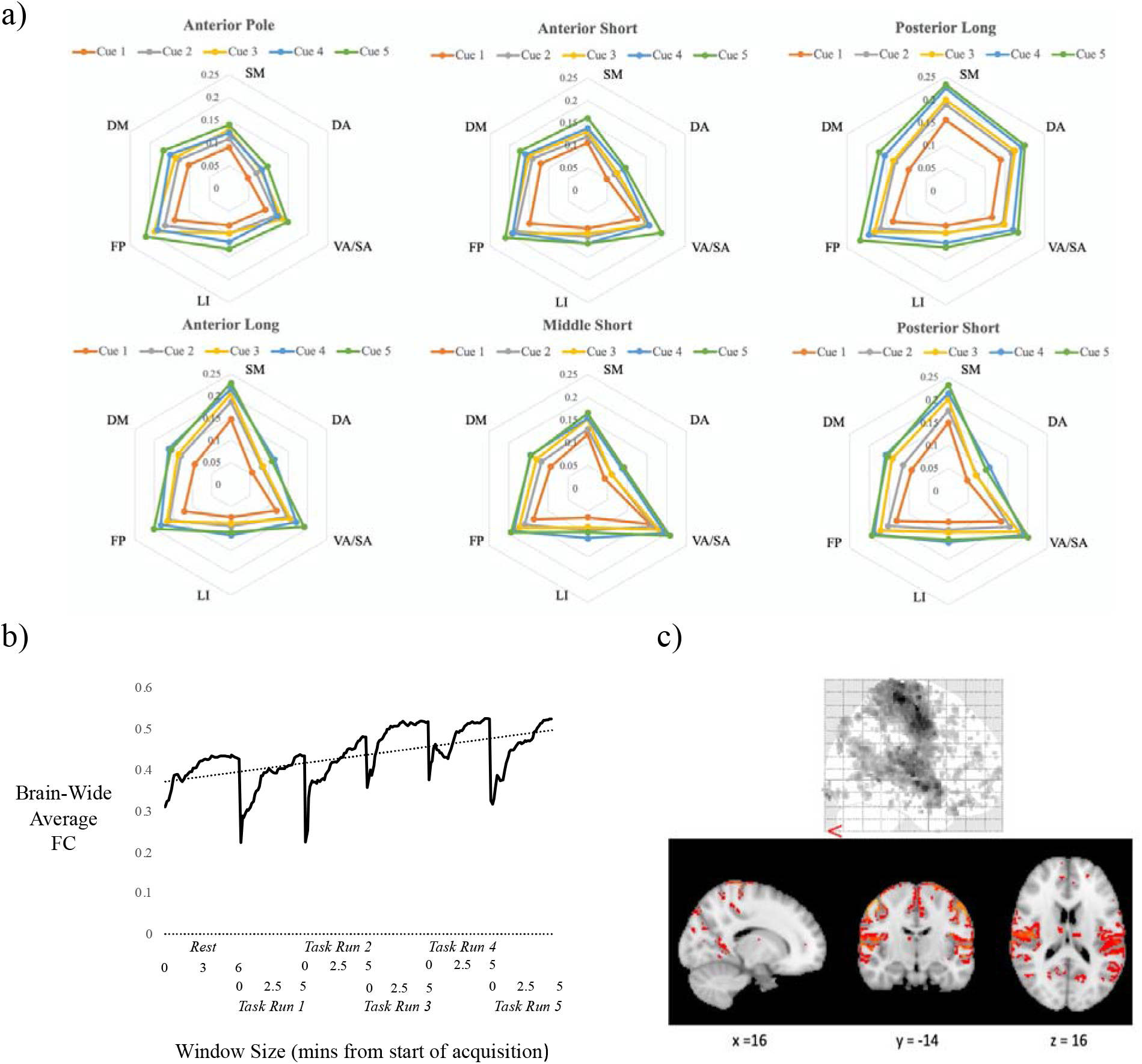
Evidence for the presence of brain-wide, spatially heterogeneous FC inflation over time in the MIC dataset. **a)** Group-average connectivity strengths (*z*-scores) in task-relevant corticostriatal circuits (i.e., between insular and striatal subregions) during five successive runs of a cue reactivity task (SM=striatal somatomotor network; DA=striatal dorsal attention network; VA/SA=striatal ventral attention/salience network; LI=striatal limbic network; FP=striatal frontoparietal network; DM=striatal default mode network). Data reveal spatially and functionally non-specific FC increases in a run-over-run manner. **b)** Expanding the scope of analysis to the whole brain. Data reveal that time-dependent FC increases are brain-wide, and occur not only during and across task acquisitions, but also during resting-state acquisitions. **c)** Voxel-wise magnitudes of FC inflation between the start and end of the task. Data reveal that FC inflation has structured spatial heterogeneity, with peak inflation rates for connections that involve the occipital lobe, sensory-motor strip and superior temporal lobe.

Surprised by the widespread, spatially non-specific nature of the time-dependent FC increases in these corticostriatal circuits, we subsequently conducted whole-brain analyses to examine whether similar time-dependent FC increases were present between other brain areas. First, we used the 100 parcel Schaefer atlas^27^ to compute a 100×100 FC matrix for each 30TR (21.6 secs) window across the task for each subject, and computed the average FC across all connections and all subjects at each window. We found that average brain-wide FC increased strongly with time within each of the 5 task runs (**Fig. 1b**; Run 1: r=0.63, *p*=0.02; Run 2: r=0.80, *p*=0.001; Run 3: r=0.49, *p*=0.07; Run 4: r=0.52, *p*=0.06; Run 5: r=0.78, *p*=0.001). Second, we used AFNI’s 3dDegreeCentrality function to quantify the average strength of each voxel’s connections with all other voxels in the brain, and evaluated how the magnitude of this quantity changed across task runs (i.e., between the first half and the second half of the task). This voxel-wise analysis revealed that the most significant (*pFWE*<0.05) time-dependent FC increases involved nodes in the sensory-motor strip, superior temporal lobe, and parts of the occipital lobe (**Fig. 1c**). These regions are not typically implicated in the cue reactivity task^28^, suggesting the presence of a non-task-related phenomenon.

To test this, we repeated the analyses for each subject’s resting-state scan. Consistent with the task data, average brain-wide FC during resting-state significantly increased with time (r=0.52, *p*=0.04; **Fig. 1b**) – from 0.33 in the first 30TR window to 0.46 by the last 30TR window (a 39% increase). Moreover, the spatial distribution of inflation magnitudes significantly corresponded to that observed during the 5 task runs (r=0.36, *p*<0.00001). Collectively, these findings suggested that the observed within-acquisition and across-acquisition brain-wide FC increases were not produced by engagement in the task or by spontaneous activity during rest, but were a spatially structured, brain-wide phenomenon linked to the duration of time spent in the fMRI scanner. We next sought to determine whether this phenomenon was also present in other, independent fMRI datasets.

### Replication of Functional Connectivity Inflation in the Human Connectome Project

Repeating the whole-brain temporal FC analyses in the HCP dataset, we replicated the MIC dataset findings of both within-acquisition and across-acquisition FC inflation. Within each of the 4, 14.4-minute HCP resting-state acquisitions, brain-wide FC increased by an average of 71%, from an average of 0.22 in the first 60TR window to an average of 0.39 by the last 60TR window (a rate of 4.9 percentage points per minute; **Fig. 2a**; compare to a rate of 6.6 percentage points per minute in the MIC dataset). During each of the 4 runs, the correlation between time and brain-wide FC was r>0.97, *p*<0.00001. Moreover, in addition to the within acquisition inflation, we found that, consistent with our results in the Discovery sample, FC inflation also occurred across sequential acquisitions in the same scanning session. In the REST1 session (which represents the first resting acquisition in the HCP data set), with few exceptions a 15-minute RL-phase encoding acquisition took place followed shortly thereafter by a 15-minute LR-phase encoding acquisition. Conversely, in the REST2 (the second resting acquisition in the HCP data set) session a LR acquisition was followed shortly thereafter by a RL acquisition. This allowed a test of whether brain-wide FC was higher in the second acquisition of a session regardless of the phase encoding direction. For both the REST1 and REST2 sessions, group brain-wide FC was significantly higher in the second acquisition of the session compared to the first acquisition at all time window durations by an average of *z*=0.04, *p*<0.00001 (**Fig. 2a**).

**Figure 2.**
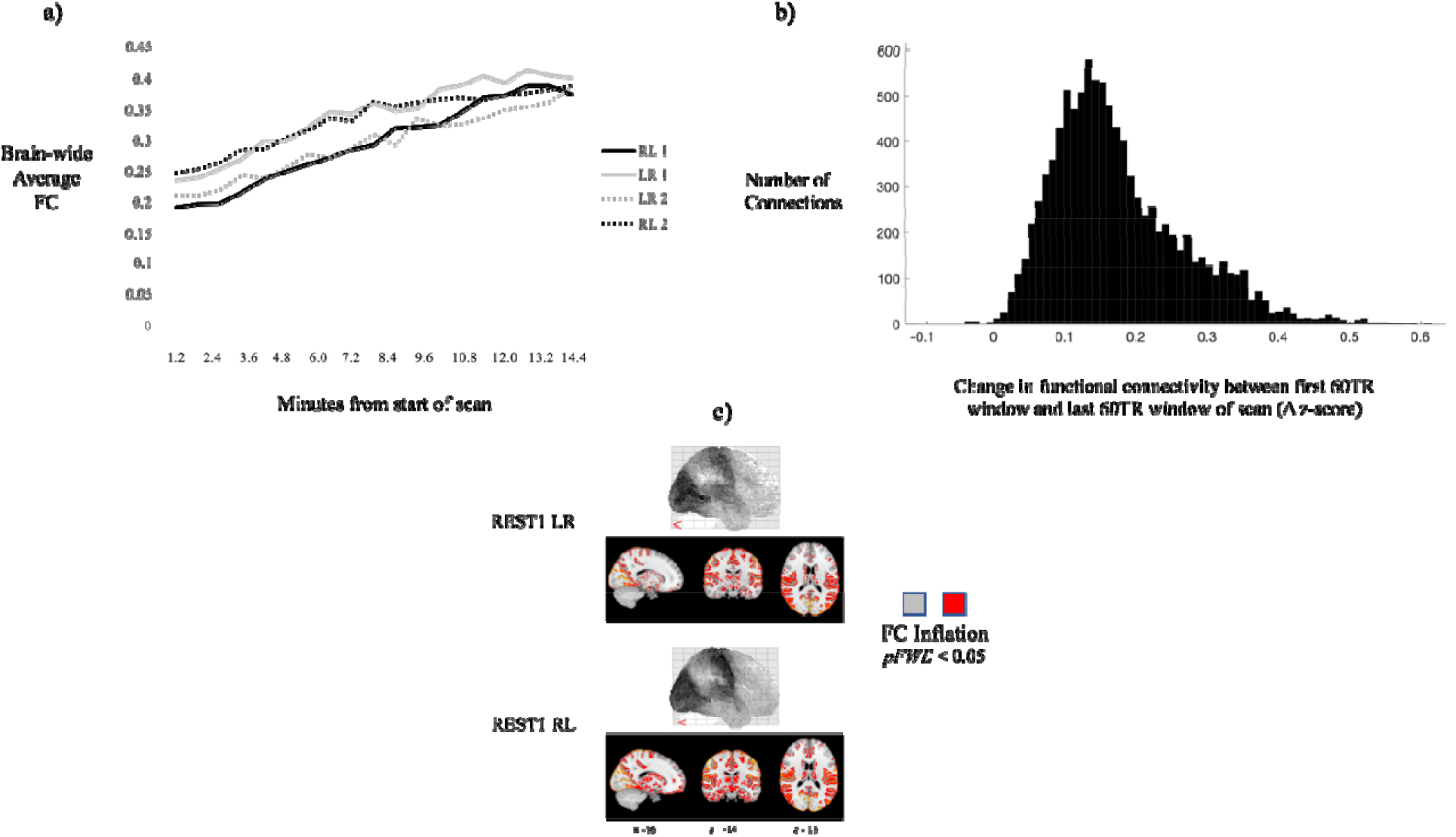
Replication of the temporal and spatial features of FC inflation in the HCP dataset. **a)** Group-average, mean brain-wide FC magnitude (*z*-score) increased monotonically with time during each of the four HCP resting-state acquisitions. Furthermore, FC magnitude increased between the first and second resting-state acquisition of each session. **b)** For a representative acquisition (REST1 LR), the distribution of FC inflation magnitudes across all connections of the connectivity matrix. **c)** Spatial visualization of FC inflation magnitudes (average FC inflation of each voxel’s set of connections). This illustrates the presence of significant FC inflation (demarcated in grayscale on surface renderings (tops) and demarcated in red on volumetric renderings (bottoms)) across the whole brain, with peaks in connections involving the occipital lobe, sensory-motor strip and superior temporal lobe.

More than 99% of examined connections displayed FC inflation. However, there was considerable spatial heterogeneity in magnitude - ranging from near Δ*z*=0 for some connections to as much as Δ*z*=0.6 for others (**Fig. 2b**). The most substantial FC inflation involved connections with nodes in the sensory-motor strip, superior temporal cortex and occipital lobe (**Fig. 2c**) - replicating the spatial pattern of FC inflation magnitude from the MIC dataset (HCP-rest and MIC rest: r=0.51, *p*<0.00001; HCP-rest and MIC-task: r=0.42, *p*<0.00001) (**Fig. 1c**).

### Functional Connectivity Inflation is Not a Consequence of ICA Denoising

To begin investigating the mechanism driving FC inflation, we first examined whether it was a computational consequence of ICA-based denoising. To do so, we repeated the above HCP analyses using HCP data with varying methods and degrees of preprocessing and denoising applied (**Fig. 3**). FC inflation was not observable in minimally preprocessed^29^ HCP data. Nor was it observable in minimally preprocessed HCP data that was denoised via motion censoring, regression of motion timeseries and regression of white matter and cerebrospinal fluid timeseries. However, it was observable in minimally preprocessed HCP data that had motion censoring and motion timeseries regression applied but did *not* have white matter and cerebrospinal fluid timeseries regression applied. Together, these data indicate that FC inflation is not an artifact of ICA-based denoising, but rather is inherent in minimally preprocessed fMRI data and becomes observable after removal of obscuring motion confounders. It also indicates that regression of white matter and cerebrospinal fluid signals - a process that is not explicitly included in ICA-based denoising - helps to mitigate FC inflation, but also introduces a considerable FC deflation trend during the first 4 minutes of scanning. Therefore, ICA-based denoising does not produce FC inflation, but rather, allows it to manifest via its ability to remove motion confounders but inability to explicitly remove white matter and cerebrospinal fluid signals. Notably, global signal regression^30^ also did not mitigate FC inflation.

**Figure 3.**
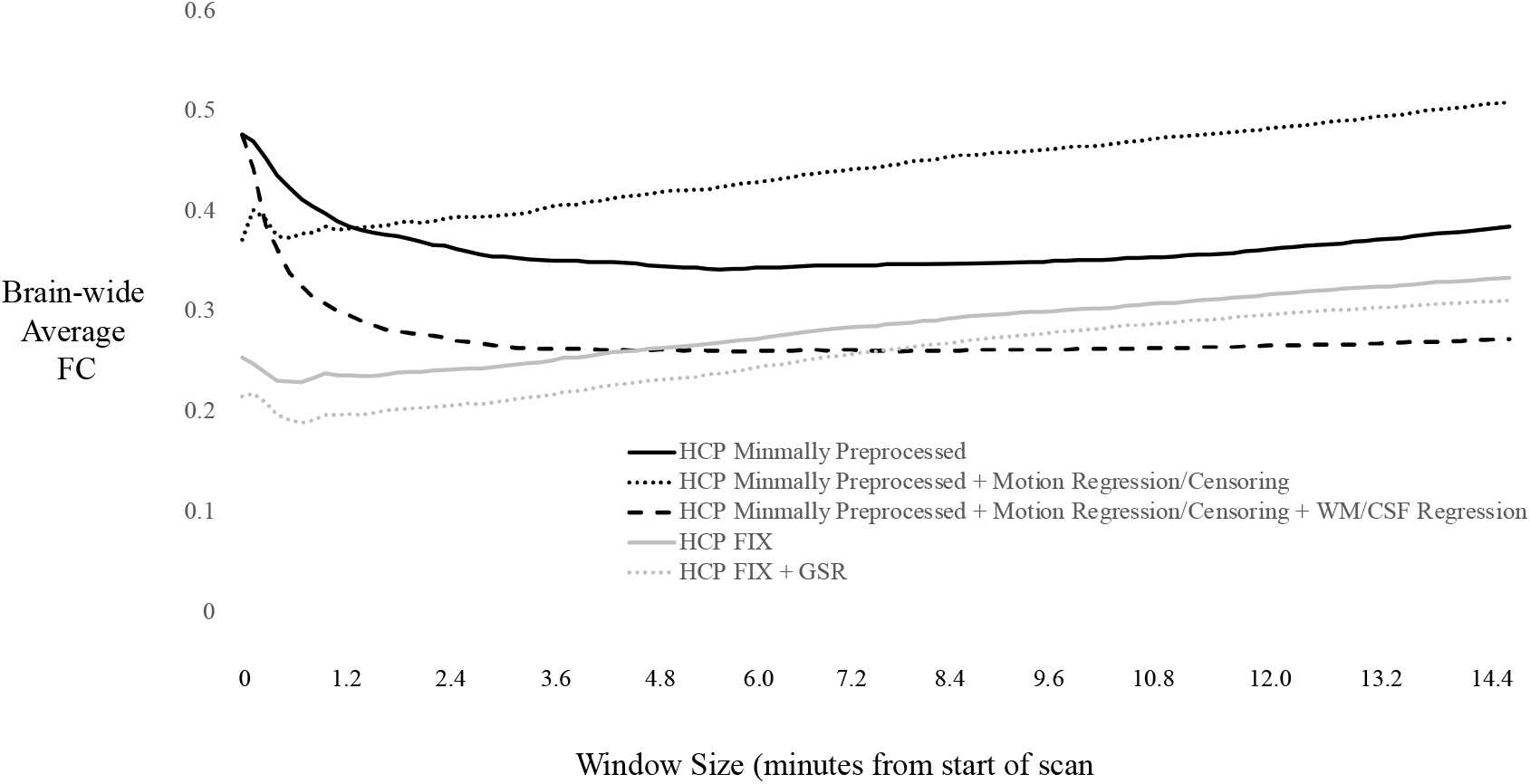
Group-averaged, mean brain-wide FC (*z*-score) computed over successively larger time windows for REST1 LR data with different preprocessing and denoising steps applied. FC inflation is present in both FIX-denoised and non-FIX denoised HCP data. FC inflation becomes apparent in minimally preprocessed data after removal of confounding motion artifacts. Neither FIX denoising nor global signal regression mitigates FC inflation.

### Functional Connectivity Inflation is Not a Consequence of Accumulating Cognitive Engagement or Habituation

To examine the possibility that FC inflation might be a non-artifactual, neurally-driven process related to habituation or extended engagement with thoughts or tasks inside the scanner, we examined whether FC inflation takes place during sleep. In the PSU dataset, average brain-wide FC computed on successive 30TR (i.e., 63 secs) windows was strongly correlated with time both in subjects’ resting-state scan (r=0.61, *p*=0.08) and sleeping-state scan (r=0.77, *p*=0.0013) (**Fig. S1**). Moreover, the spatial pattern of FC inflation magnitude in the resting-state and sleep-state scans corresponded strongly with each other (r=0.50, *p*<0.00001) and with the resting-state and task patterns from the other independent datasets (MIC-rest and PSU-rest: r=0.39, *p*<0.00001; MIC-task and PSU-rest: r=0.37, *p*<0.00001; MIC-rest and PSU-sleep: r=0.38, *p*<0.00001; HCP-rest and PSU-sleep: r=0.40, *p*<0.00001). The ubiquitous presence of corresponding spatial maps of FC inflation across rest, task and even sleep conditions indicates that it is not the consequence of habituation to or accumulating engagement with thoughts or stimuli in the scanner.

### FC Inflation is Temporally Paralleled by Inflation of Global Mean Signal (GMS) Variance

Having provided evidence that FC inflation is not a computational consequence of ICA denoising and is not driven by a neurally-based process, we next turned to possible non-neural, physiological noise sources. If the source of FC inflation is noise-based, we would expect to observe an increase in noise over time in the low-frequency band (i.e., in the frequency band of neural activity). To obtain a measure of relative systemic noise power over time, we computed the time course of each HCP subject’s GMS (i.e., the sum of the low frequency BOLD signals throughout the brain at each time point) in each resting scan, normalized each time course by its mean (i.e., converted the signal to percent BOLD change), and calculated the variance at every time point across all subjects for each scan. In doing so, we found that systemic low-frequency noise power - measured by the variance of the normalized GMS at each time point – continuously increased over time both within individual scans and across scans in the same session (**Fig. 4**). This strongly paralleled the temporal features of FC inflation in this dataset (**Fig 2a.)**. Collectively, this data suggested that a time-dependent, low-frequency, non-neuronal noise signal may underlie the temporal inflation of FC and GMS variance in the scanner.

**Figure 4.**
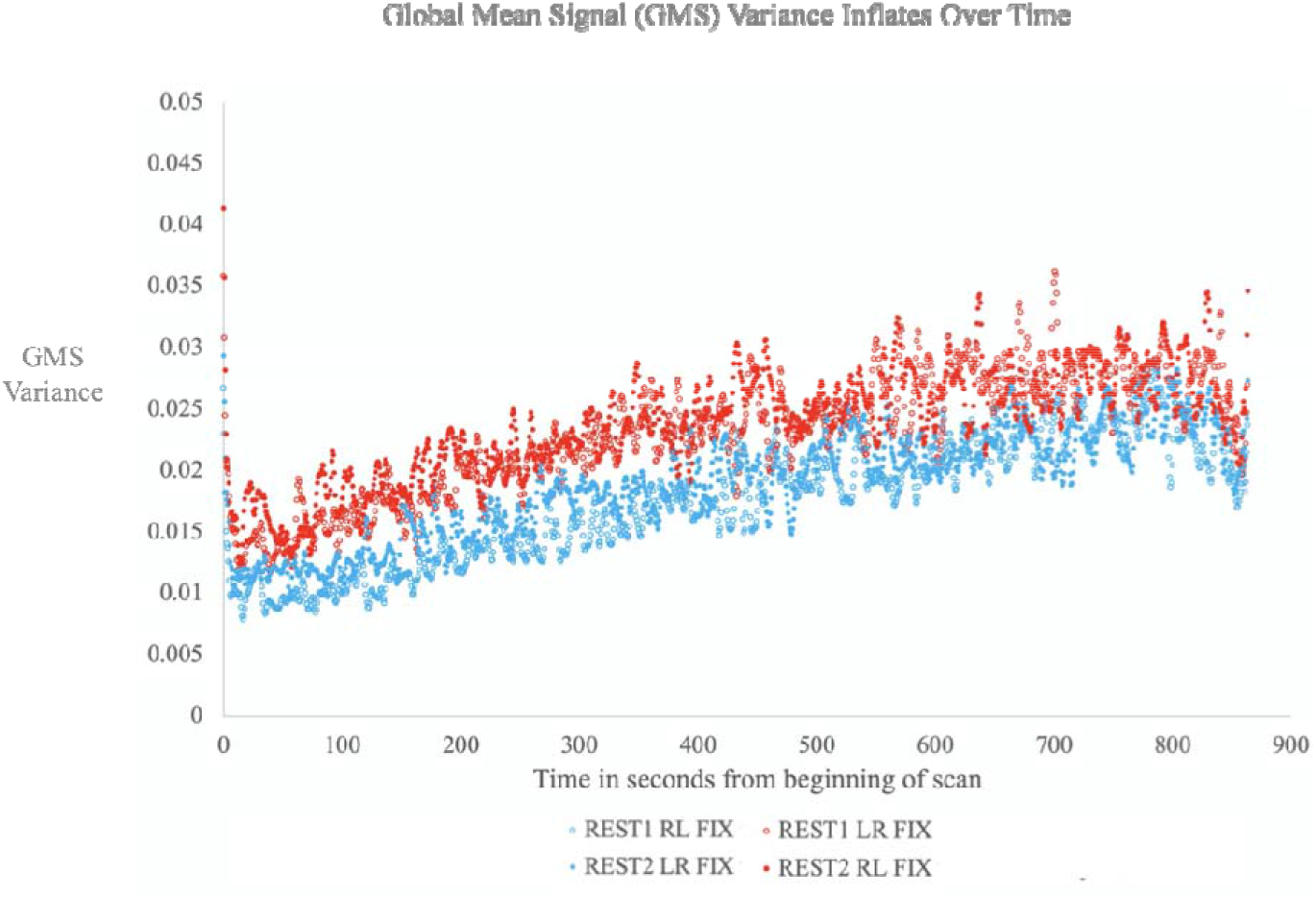
GMS variance inflates over time during and across scanning acquisitions. Graph displays percent normalized GMS variance in the low frequency band (0.01-0.15Hz) across all subjects, for each of the HCP REST1 and REST2 acquisitions after ICA-FIX denoising. The variance grows monotonically with time within each acquisition, and the second acquisition of each session (i.e., LR for REST1 and RL for REST2, denoted with red markers) starts and finishes with a higher variance than the first acquisition of each session (i.e., RL for REST 1 and LR for REST2, denoted with blue markers).

### Temporal Inflation of the Systemic Low Frequency Oscillation (sLFO) Signal Drives GMS Variance Inflation and FC Inflation

We next sought to determine the particular non-neuronal noise source driving these time-dependent effects. Multiple non-neuronal noise sources are known to contribute to BOLD signal variability, including subject head motion, scanner and acquisition artifacts, and physiological processes such as cardiac and respiratory cycles. However, these noise signals are known to be captured and filtered out of fMRI data by the spatial ICA denoising methods used in the examined datasets (e.g., ICA FIX)^31^. Moreover, the spatial distributions of these noise signals do not align with the spatial distribution of the noise signal under examination here (**Fig. 1c, Fig. 2c**). Given these two constraints (i.e., a signal that would not be removed by ICA FIX and that displays the spatial distribution observed here), a strong candidate was the systemic low frequency oscillation (sLFO) signal.

The sLFO is a recently identified non-neuronal, spontaneous physiological oscillation that travels with the blood and is thought to have an extracerebral origin^4,32-34^. A key feature of the sLFO signal that distinguishes it from other low-frequency physiological signals that are captured by spatial ICA denoising is its dynamic propagation across the brain. The sLFO displays a pattern of time delayed appearances across different regions of the brain, consistent with delays that would be expected if the signal were moving through the brain’s vasculature at the rate of cerebral blood flow^4^. Spatial ICA denoising methods like ICA-AROMA and ICA-FIX are not well suited to capturing noise signals that appear in different brain regions at different times, and thus the sLFO signal is one that would evade detection and removal by these denoising methods. Moreover, spatial maps specifying the contribution of the sLFO to BOLD signal variance in different areas of the brain align with the spatial distribution of FC inflation observed here, with peaks in the occipital lobe, sensory-motor strip and superior temporal lobe^32,35,36^. The sLFO signal would thus satisfy the constraints specified above.

However, the sLFO signal is not previously known to increase in strength over time in the scanner – a final necessary feature if it is to account for the temporal increases in the GMS variance and FC magnitude observed here. As such, we investigated the temporal evolution of sLFO signal strength during fMRI scanning in the HCP resting-state acquisitions. To isolate and measure the sLFO signal, we used Regressor Interpolation at Progressive Time Delays (RIPTiDe) as implemented by the software package rapidtide^37,38^, which has been used extensively to both characterize and remove physiologically based sLFO signals^32,39-44^

The RIPTiDe analysis revealed that the proportion of BOLD signal variance attributable to sLFO noise increased monotonically with time within both the HCP’s first and second resting state acquisitions of a given scanning session, and also increased across these acquisitions (**Fig. 5a**). These temporal properties matched those of both the observed GMS variance inflation (**Fig. 4**) and FC inflation (**Fig. 2a**). Moreover, the voxel-wise spatial distribution of sLFO signal inflation magnitude (**Fig. 5b**) corresponded strongly to the voxel-wise spatial distribution of FC inflation magnitude (RL 1: r=0.64, *p*<0.00001; LR 1: r=0.63, *p*<0.0001). Just as in the FC inflation maps, sLFO signal inflation was greatest in the occipital lobe, sensory-motor strip, and superior temporal lobe. In these regions, up to 45% of low-frequency BOLD variance was attributable to sLFO noise by the final quarter of the acquisition (**Fig. 5b**). Matching spatiotemporal patterns of sLFO signal inflation were also present in both the MIC rest and task data (**Fig. S2a-b**) and the PSU rest and sleep data (**Fig. S2c-d**), indicating the ubiquity of this physiological phenomenon across fMRI datasets.

**Figure 5.**
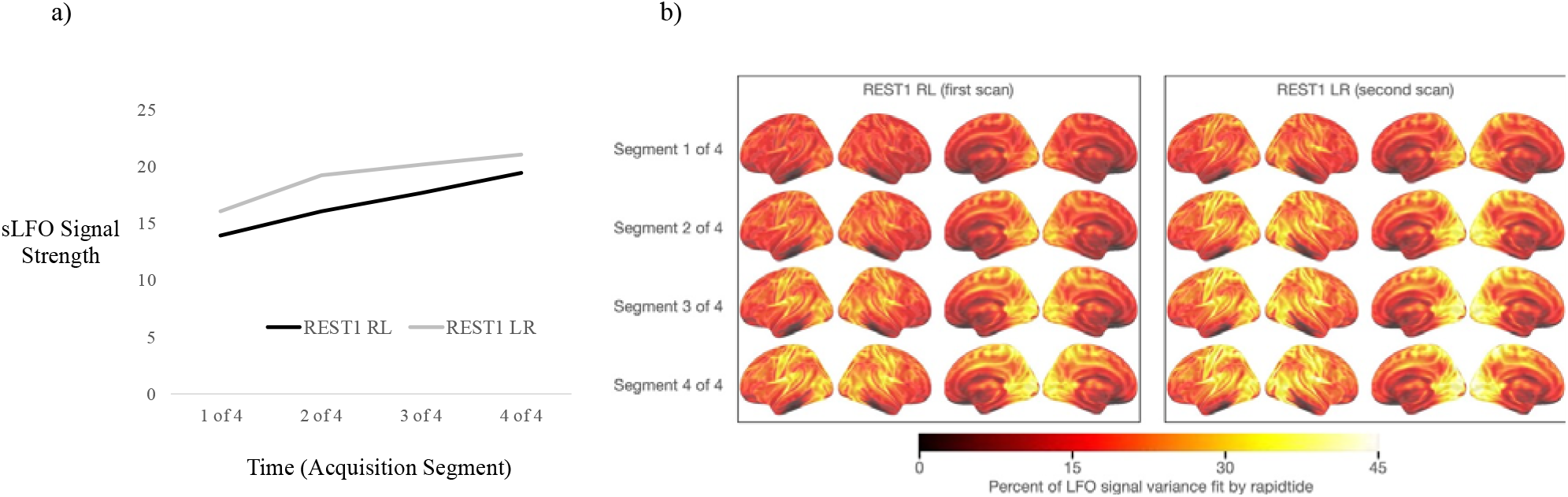
Temporal and spatial properties of the sLFO signal in HCP REST1 data. **a)** Brain-wide average percent of low frequency (0.01-0.15Hz) BOLD variance that was attributable to the non-neuronal sLFO signal at sequential time points (i.e., each quarter of an acquisition) in the HCP REST1 data. The degree of sLFO noise detected increased monotonically in both the first (RL) and second (LR) acquisitions across time segments; furthermore, the second acquisition (LR) had a larger sLFO contribution than the first. This matched the increase over time in both global mean signal variance and functional connectivity strength during these acquisitions (see **Fig. 2a** and **Fig. 4**). **b)** Voxel-wise spatial distribution of the percent of low-frequency BOLD variance attributable to the sLFO signal during each quarter of the REST1 RL and LR acquisitions. These spatial maps corresponded strongly to the spatial maps of FC inflation (RL 1: r=0.64, *p*<0.00001; LR 1: r=0.63, *p*<0.0001).

### Removal of the sLFO Signal via RIPTiDe Mitigates Global Mean Signal (GMS) Variance Inflation and FC Inflation

Finally, to solidify the causal role of the sLFO signal in driving FC inflation, and to evaluate a practical procedure for mitigating FC inflation in fMRI datasets, we examined the impact of sLFO signal denoising via RIPTiDe on FC inflation.

In the HCP resting-state acquisitions, RIPTiDe significantly attenuated the inflation of global mean signal variance over time, producing a small increase over time relative to the average value (**Fig. 6a**). Accordingly, RIPTiDe also reduced brain-wide FC inflation between the start and end of acquisitions from 71% (before RIPTiDe) to 27% (after RIPTiDe), reducing the range of average brain-wide FC magnitudes from 0.22 to 0.39 (before RIPTiDe) to 0.16 to 0.21 (after RIPTiDe) (**Fig. 6b**). On a regional voxel-wise basis, RIPTiDe eliminated FC inflation in the frontal cortex and in much of the parietal and temporal lobes (**Fig. 6c**). It significantly reduced, though did not fully eliminate, FC inflation in the occipital lobe, sensory-motor strip and superior temporal lobe. Overall, RIPTiDe significantly (p<0.000001) reduced the average magnitude of FC inflation experienced by individual brain connections between the start and end of HCP resting-state acquisitions from 0.17 (before RIPTiDe) to 0.03 (after RIPTiDe) (**Fig. 6d**). Complementing FIX ICA denoising with RIPTiDe denoising of the sLFO signal optimized within-acquisition stability of brain-wide FC better than other denoising procedures, including regression of WM and CSF (**Fig. S3**).

**Figure 6.**
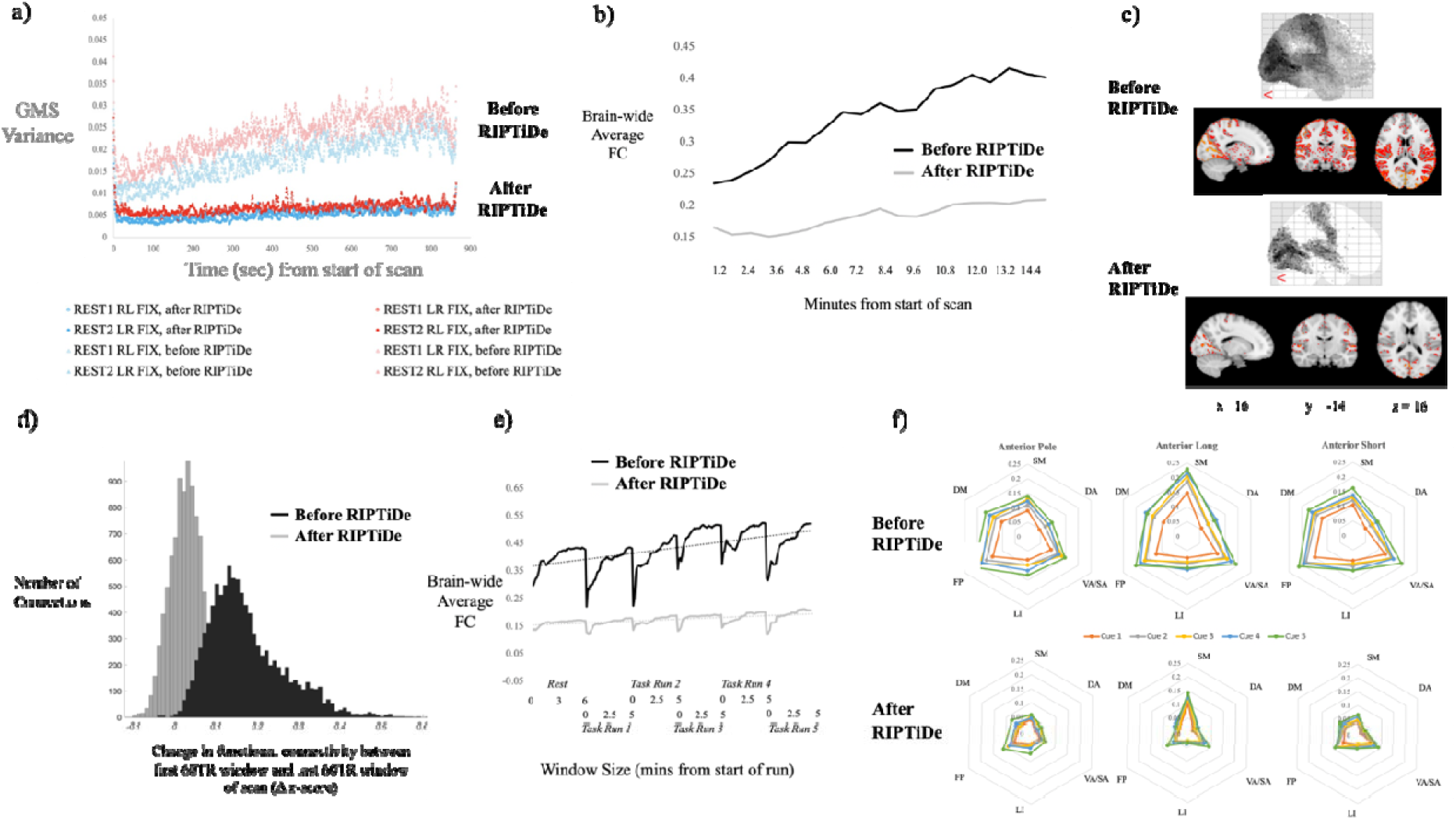
The impact of RIPTiDe denoising. Removing the sLFO signal from HCP FIX rs-fMRI data via RIPTiDe substantially attenuated both GMS variance inflation (a) and FC inflation (b). It eliminated statistically significant FC inflation throughout the brain except in peak sLFO inflation regions (occipital lobe, sensory-motor strip, and superior temporal lobe) (c), and reduced the average magnitude of FC inflation for all connections from 0.17 to 0.03 (d). Similarly, in the MIC dataset, RIPTiDe denoising eliminated significant within-run and across-run FC inflation during the cue reactivity task (e), and revealed that pre-RIPTiDe findings of apparent task-associated FC increases in insula-striatal circuits were false positive results driven by artifactual FC inflation (f).

In the MIC dataset, removal of the sLFO signal via RIPTiDe significantly reduced both within-acquisition and across-acquisition FC inflation (**Fig. 6e**). Across all rest and task acquisitions, RIPTiDe denoising reduced FC inflation between the beginning and ends of acquisitions from an average of 46% (before RIPTiDe) to an average of 23% (after RIPTiDe), reducing the range of average brain-wide FC magnitudes from 0.34 to 0.49 (before RIPTiDe) to 0.17 to 0.20 (after RIPTiDe). In doing so, RIPTiDe denoising eliminated all significant voxel-wise FC increases initially found between the first half and second half of the task (**Fig. 2c**), and rendered null our initial significant findings of apparent task-related, run-over-run increases in insula-striatal connectivity (**Fig. 6f**).

Collectively, these data provide strong evidence that the non-neuronal sLFO signal is the causal driver of FC inflation (and as a side effect, the inflation of the GMS). The sLFO signal’s temporal and spatial features correspond significantly to those of FC inflation, and once the sLFO signal is removed from the data, FC inflation is substantially reduced.

It is important to note that while the GMS and the sLFO signal are closely related, they are not interchangeable. The GMS is largely an epiphenomenon of averaging the sLFO signal over a spatially complex pattern of time delays. Regressing out the sLFO (with voxel specific delays applied) will effectively attenuate the GMS, but the converse is not true. Regressing out the GMS from every voxel does not account for the delay structure of the sLFO signal throughout the brain, and will therefore incompletely remove the sLFO (and the resulting artifactual connectivity). We demonstrate the inability of GMS regression to attenuate FC inflation in **Fig. 3**. Moreover, GMS regression without accounting for delay is the cause of the artifactual spurious negative correlations that are a well-known side effect of the technique^32^.

### FC Inflation Has Negligible Impact on Between-Scan Reproducibility and Subject Discriminability

As a post-hoc analysis, we sought to understand how FC inflation may have been overlooked in prior investigations of fMRI BOLD signal reliability, which have principally focused on between-scan reproducibility and individual discriminability. The present data indicate that FC inflation’s considerable impact on within-scan reliability and spatiotemporal validity does not extend to between-scan reproducibility and individual discriminability. For example, in the HCP dataset, even as FC inflation increasingly compromised within-scan reliability throughout each 15-minute acquisition, between-acquisition reproducibility remained high (94-98%), plateauing after 7 minutes of scanning (**Fig. S4a**). In addition, substantial RIPTiDe-driven improvements to within-scan reliability (an 81.2% reduction in FC inflation) produced only small changes in between-scan reproducibility (6.5% increase) (**Fig. S4b)** and ICC-indexed subject discriminability (7.4% decrease) (**Fig. S4c**). This is consistent with prior reports showing that while physiological denoising methods increase the validity of fMRI data and increase between-scan reproducibility, they tend to decrease subject discriminability^39,45^ via the removal of ICC-inflating noise. The relative insensitivity of between-scan reproducibility and subject discriminability to FC inflation can be understood when considering that these metrics use time-averaged connectivity values as inputs, which would obscure the impact of temporal changes in connectivity that happen within a scan.

## Discussion

The presence of the non-neuronal sLFO signal, its sizable contribution to BOLD signal variance, and its potential for impacting the validity of functional connectivity measurements has been appreciated for over a decade^32,38,46,47^. However, core features of the sLFO signal – that it occupies the same low-frequency band as neural functional connectivity and manifests across the brain in a time-delayed manner – make it particularly challenging to capture and regress out of fMRI BOLD data. Moreover, despite recognition of the sLFO signal’s effect on the validity of functional connectivity estimates^47^, prior work has not indicated that it drives substantial spatial or temporal distortions. Likely for these reasons, sLFO denoising has not been adopted as a step in field-standard fMRI preprocessing and denoising pipelines.

Crucially, however, it was not previously recognized that the sLFO signal undergoes substantial time-dependent inflation during fMRI scanning, and that this drives significant temporal inflation of both GMS variability and brain-wide FC magnitude across rest, task, and sleep conditions. Hints of time-dependent increases in the variability of the GMS^48^ and BOLD signal amplitude^49^ during resting state scans in circumscribed brain networks are evident in the literature. However, the implications of these phenomena for the within-scan stability of FC magnitudes have not been previously identified, and potential underlying mechanisms previously examined (i.e., temporal increases in subject head motion^49^ and breathing rate variability^48^ during scanning) were ultimately found to be largely independent and correlational only. Here, by demonstrating the tight correspondence between the non-neuronal sLFO signal–s spatiotemporal features and those of FC inflation, coupled with the substantial reduction in GMS variability and FC inflation that occurs when the sLFO signal is removed from the data, we strongly implicate the sLFO signal as the causal driver of artifactual FC inflation in fMRI datasets – and, in doing so, point to a direct FC inflation mitigation strategy.

sLFO signal inflation appears to be ubiquitous across human fMRI datasets, including those acquired with both slow TRs (e.g., 2.1 sec) and fast TRs (e.g., 0.72 sec), and including all types of scanning conditions (i.e., rest, task, and sleep). We have demonstrated here how sLFO-driven FC inflation can produce false positive findings, suggesting that the results of many prior studies examining FC may be distorted by FC inflation. However, the impact of sLFO signal inflation on prior investigations of FC depends both on the denoising methods used and the types of analyses conducted. Datasets using ICA-based denoising are the most highly susceptible to sLFO-driven FC inflation. This is concerning given that the HCP’s ICA-FIX dataset is among the most widely used analysis-ready resting-state datasets in the field. That said, the manifestation of sLFO-driven FC inflation in ICA denoised data is a testament to ICA’s efficacy in removing other confounding noise that obscures the sLFO effect. Conversely, datasets using a denoising procedure that includes regression of white matter and cerebrospinal fluid timeseries attenuate FC inflation considerably relative to ICA denoised data. However, while regressing out white matter and cerebrospinal fluid signals (which are highly correlated with the GMS at different time delays^47,50^) performs a crude approximation of RIPTiDe and will remove some fraction of the sLFO signal, this technique is not an adequate substitute for the voxel specific sLFO regression applied by RIPTiDe.

For those datasets strongly impacted by FC inflation, analyses that investigate FC temporal dynamics – changes in FC over time during scanning – would be most affected. These include analyses assessing changes within individual acquisitions and across acquisitions from the same scanning session. For instance, “dynamic connectivity” analyses seek to understand how patterns of FC change and cycle through different configurations within individual acquisitions. Since FC inflation causes the FC strength of certain connections to artificially increase at a faster rate than others, this could meaningfully distort the makeup and spatiotemporal properties of observed configurations. Similarly, analyses of FC changes during and across task run acquisitions would be liable to conflate artificial, sLFO-driven FC increases with task effects. Moreover, FC inflation and its structured spatial heterogeneity would distort attempts to chart accurate normative brain connectivity maps.

On the other hand, FC inflation would likely have minimal, if any, effect on other types of analyses. For instance, comparisons between groups within the same dataset would be negligibly impacted by FC inflation – as long as both groups experience similar sLFO signal inflation in the scanner. This may not be the case, however, for comparisons between healthy groups and patient groups affected by neurovascular deficits (e.g., Alzheimer’s Disease) where blood flow-driven sLFO dynamics themselves may differ between groups. In addition, within-subject longitudinal analyses, where FC is compared between scans acquired during independent scanning sessions (e.g., on different days) would be negligibly affected by FC inflation, as we would expect similar magnitudes and patterns of FC inflation to occur in the same subject during both scans. It is worth noting, however, that in these cases, while the inflated noise may not be different between conditions, it is still present. Thus, instead of comparing neuronal connectivity alone, there is also an implicit comparison of the degree of artifactual physiological connectivity, which may be affected by state or group differences in brain circulatory patterns.

It is noteworthy that burgeoning fMRI experimental design improvements (e.g., longer scan durations and larger sample sizes), which have improved between-scan reproducibility and subject discriminability, do not address FC inflation – a problem of “within-scan” reliability and spatiotemporal validity. The nature and origin of FC inflation help to explain why. While data aggregation approaches (e.g., longer scan durations) help to cancel out the random noise that degrades between-scan reproducibility and subject discriminability, they solidify and amplify structured noise of the type that drives FC inflation. Indeed, perhaps ironically, longer scan durations actually produce more spatiotemporally distorted FC maps than shorter scans.

Importantly, however, we demonstrate here that FC inflation can be easily addressed by implementing RIPTiDe – a publicly available sLFO signal denoising procedure – as a standard step in fMRI preprocessing and denoising pipelines. We show that RIPTiDe can be used in conjunction with ICA to provide the benefits of ICA denoising while also attenuating FC inflation. Notably, RIPTiDe considerably reduces, but does not completely eliminate, FC inflation. This suggests that a small component of FC inflation may be driven by a non-sLFO source, such as a scanner artifact.

An important direction in subsequent work will be to ascertain why the non-neuronal sLFO signal undergoes time-dependent inflation during fMRI scanning, to understand and perhaps mitigate this phenomenon. The spatial locations of the strongest sLFO inflation magnitudes (occipital lobe, sensory-motor strip, and superior temporal lobe) underlay areas of the skull that make contact with the scanner table and head coil. This suggests that one possibility could be a buildup of cranial pressure that impacts cerebral blood flow. Another possibility could be related to physiological or hydrostatic effects resulting from subjects’ transition from an upright to a supine position in the scanner^51^. A third alternative is suggested by Aso’s recent finding that increases in sLFO and global signal intensities are significantly correlated with a decrease in respiratory and pulse rate^52^, indicating that it may be a secondary effect of decreased arousal or physiological habituation to the scan environment over time. Finally, the change in signal strength could relate to increases in local CO2 concentration around the subjects’ face in the relatively still air of the magnet bore, leading to progressively increasing vasodilation^53^. Some combination of these factors may be responsible for the continuous inflation of the LFO signal during scanning; this will be systematically examined in follow-up work.

## Methods

### Subjects

#### HCP

Data used in these analyses include fMRI scans from individuals gathered as part of the Human Connectome Project (HCP) 1200 subject release. A detailed description of the recruitment for the HCP is provided by others^20,21^. Briefly, individuals were excluded by the HCP if they reported a history of major psychiatric disorder, neurological disorder, or medical disorder known to influence brain function. We further excluded individuals from the HCP database that reported a family history of schizophrenia, met DSM-IV criteria for alcohol dependence, or reported a lifetime history of repeated substance use (>10 instances of cocaine, hallucinogen, opiate, sedatives, or stimulant use, >20 instances of tobacco use or >100 instances of marijuana use). In addition, participants included in the present analyses provided a breath sample indicating <0.05 blood alcohol content on the day of scan and a urine sample that was negative for any substances of abuse (cocaine, marijuana, opiates, amphetamine, or methamphetamine). The final data set included 462 individuals (Female = 281) who were an average age of 28.66 years of age (±3.65, range 22–36) and reported completing an average of 15.34 years of education (±1.59, range 11–17). Three hundred and fifty-one participants self-identified as White, 64 identified as Black or African American, 35 identified as Asian, Native Hawaiian, or Pacific Islander, six identified as multiracial, and six were unknown or not reported.

#### MIC

63 nicotine-dependent individuals (24 women; age 28.7±6.7; education 15.0±2.0; 47 White, 6 Black, 6 Asian, 4 multi-racial) who reported an interest in quitting smoking and participated in a fMRI scanning session that included a 6-minute resting-state acquisition and 5 runs of a cue reactivity task. A Fagerström Test for Nicotine Dependence^54^ (FTND) score ≥ 4 and an expired carbon monoxide concentration of ≥ 5 ppm at the time of screening were required. Serious medical illness, pregnancy (confirmed by urinalysis) drug or alcohol (except nicotine) dependence, major depressive disorder within the past 6 months, and current or lifetime history of schizophrenia, schizoaffective disorder, bipolar disorder, or psychotic disorders not otherwise specified (confirmed via Structured Clinical Interview for DSM (SCID) IV) were exclusionary criteria. Abstinence from drug/alcohol use was confirmed by urine/breath samples respectively (QuickTox11 Panel Drug Test Card, Branan Medical Corporation, Irvine California; Alco-Sensor IV, Intoximeters Inc., St. Louis, MO). All procedures were completed at McLean Hospital and were approved by the Partners Human Research Committee. Participants provided written and verbal informed consent after receiving a complete study description.

#### PSU

40 healthy participants were initially recruited in this experiment with provided informed written consent. Four participants did not complete the entire experimental protocol and three participants were excluded due to technical problems. Therefore, the final sample size contained a total of thirty-three participants (age: 22.1±3.2 years; male/female: 17/16). All experimental procedures were approved by the Institutional Review Board at the Pennsylvania State University. During the sleep scans, participants were instructed to try to fall asleep. If awake at any point during the sleep scan, subjects were instructed to hit a button once per second to provide a behavioral measurement of wakefulness. Participants were instructed to allow themselves to stop hitting the button as they felt themselves falling asleep. Sleep states were confirmed via simultaneous EEG recording.

### fMRI Acquisition

#### HCP

HCP neuroimaging data were acquired with a standard 32-channel head coil on a Siemens 3 T Skyra modified to achieve a maximum gradient strength of 100 mT/m^12,55,56^. Gradient-echo EPI images were acquired with the following parameters: TR□= □720 ms, TE□= □33.1 ms, flip angle□= □52°, FOV□= □280□× □180 mm, Matrix□= □140□× □90, Echo spacing□= □0.58 ms, BW□= □2290 Hz/Px. Slice thickness was set to 2.0 mm, 72 slices, 2.0 mm isotropic voxels, with a multiband acceleration factor of 8.

#### MIC

Scanning was performed on a Siemens Prisma 3T scanner (Erlagen, Germany) with a 64-channel head coil. Functional scans (resting and cue reactivity) were acquired with TR = 720 ms, TE = 30 ms, slices = 66, phase encode direction posterior to anterior, Flip angle = 66o voxel size = 2.5 × 2.5 × 2×5 mm, GRAPPA factor of 2, and a multi-band acceleration factor = 6. After all functional scans, multiecho multi-planar rapidly acquired gradient echo-structural images were acquired with the following: TR = 2530 ms, TE1= 3.3 ms, TE2 = 6.98 ms, TE3 = 8.79 ms, TE4 = 10.65 ms, flip angle 7o, resolution = 1.33 × 1 × 1 mm.

#### PSU

Blood oxygen level dependent (BOLD) functional MRI data were acquired using an echo planar imaging (EPI) sequence with the following parameters: TR = 2100 milliseconds, TE = 25 milliseconds, flip angle = 90 degrees, slice thickness = 4 mm, number of slices = 35, FOV = 240 mm, in-plane resolution = 3mm × 3mm.

### fMRI Preprocessing

#### HCP

The HCP minimal preprocessing pipeline for resting-state data removes spatial distortions, realigns volumes to compensate for subject motion, registers the echo planar functional data to the structural data, reduces the bias field, normalizes the 4D image to a global mean, and masks the data with a final FreeSurfer-generated brain mask^29^. HCP FIX data additionally applies ICA-FIX to denoise the minimally preprocessed data of motion artifacts.

#### MIC

Data were analyzed using tools from the Functional Magnetic Resonance Imaging of the Brain (FMRIB) Software Library (FSL: www.fmrib.ox.ac.uk/fsl). Standard pre-processing was conducted on both the resting and task-based data including: motion correction with MCFLIRT, brain extraction using BET, slice time correction, spatial smoothing with a Gaussian kernel of full-width half-max of 6mm, and high pass filtering at 0.01 Hz. The only non-FSL tool used was spikefix (https://github.com/bbfrederick/spikefix), which evaluates fMRI data for motion/intensity spikes. For task-based data, these spikes are removed and single point confound regressors representing noise-related timepoints were generated. Resting-state data was denoised via independent component analysis using the FSL tool for multivariate exploratory linear decomposition into independent components (MELODIC). MELODIC was run for each subject and spatial/temporal information for each independent component was visually inspected to identify noise-related components, which were then regressed out of the resting-state fMRI data to generate a denoised fMRI timeseries.

#### PSU

Preprocessing was performed using FMRIPREP version 20.2.7. Functional data was slice time corrected using 3dTshift from AFNI v16.2.07 [16, RRID:SCR_005927] and motion corrected using mcflirt (FSL v5.0.9). This was followed by co-registration to the corresponding T1w using boundary-based registration with six degrees of freedom, using bbregister (FreeSurfer v6.0.1). Motion correcting transformations, BOLD-to-T1w transformation and T1w-to-template (MNI) warp were concatenated and applied in a single step using antsApplyTransforms (ANTs v2.1.0) using Lanczos interpolation. Data was denoised via independent component analysis using the FSL tool for multivariate exploratory linear decomposition into independent components (MELODIC). MELODIC was run for each subject and spatial/temporal information for each independent component was visually inspected to identify noise-related components, which were then regressed out of the resting-state fMRI data to generate a denoised fMRI timeseries.

### Regressor Interpolation at Progressive Time Delays (RIPTiDe)

To estimate the physiological noise contamination in each dataset, we performed “Regressor Interpolation at Progressive Time Delays” (RIPTiDe) analyses^32^ using “rapidtide”, a performance optimized open source implementation of the RIPTiDe processing stream (https://github.com/bbfrederick/rapidtide)^57^, to generate whole-brain blood arrival times. The RIPTiDe procedure has been described in detail previously, and involves finding the sLFO signal in minimally preprocessed data, and then removing it from denoised (e.g., via ICA) data^4^. Briefly, the rs-fMRI time courses in each voxel is filtered to the LFO band (0.009 – 0.15 Hz). Then, a “reference regressor” representing an estimate of the “systemic LFO” (sLFO) signal, is extracted and refined from the LFO-filtered global mean BOLD signal using an iterative “bootstrapping” process by successively calculating the arrival time of the reference regressor in each voxel using cross-correlation, time shifting each voxel’s time course to align the sLFO component, and summing the aligned time courses to generate a new, “sharpened” estimate of the reference regressor^45^. The estimation procedure generally converges after 3 steps, and the final cross-correlation between the refined reference regressor and the LFO-filtered rs-fMRI time course is used to determine sLFO arrival time in each voxel. The peak of the cross-correlation in each voxel (within a time window set between -10 to 10 seconds) is fit with a Gaussian function to determine the time delay and the maximum correlation coefficient^58^. The maximum correlation coefficient, R, represents the strength of the sLFO contribution to the rs-fMRI BOLD signal in a given voxel (100 * R^2 is the percentage of variance in the voxel explained by the time-shifted sLFO regressor), and the blood arrival time is defined as the time delay at which the correlation between a given voxel’s time course and the reference regressor is maximal. In the final step, voxel specific regressors are generated by time shifting the refined sLFO regressor by the blood arrival time in each voxel, and a GLM filter is used to fit and remove the signal from the fMRI data. As we showed previously, this removes physiological contamination without producing spurious negative correlations between regions^32^.

## Acknowledgements/Funding

This work was supported by 5R01DA039135-06 (CK), the Intramural Research Program of the NIH, NIDA (ACJ), and 1RF1MH130637-01 and 1R21AG070383-01 (BBF).

## Data Availability

The HCP dataset is publicly available on an open access repository (https://db.humanconnectome.org/app/template/Login.vm), which can be accessed after signing a data use agreement. The PSU dataset is publicly available on the OpenNeuro repository (https://openneuro.org/datasets/ds003768/versions/1.0.9). The MIC dataset is available upon reasonable request to the authors.

## Code Availability

Code for performing RIPTiDe denoising via the rapidtide package is publicly available (https://github.com/bbfrederick/rapidtide).

