## Supplement for "Brain-wide functional connectivity artifactually inflates throughout fMRI scans: a problem and solution"

**Supplemental Information**

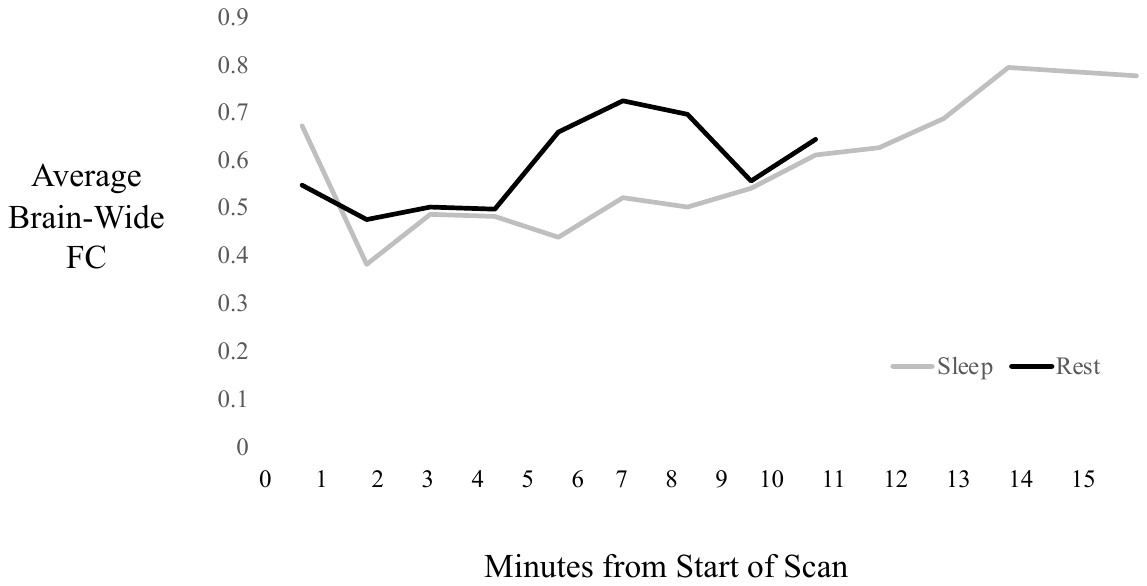

**Figure S1.** FC inflation is present even during sleep-state acquisitions. Group-average, mean brain-wide FC strength (*z*-score) at successive 30 TR time windows (i.e., every 63 secs) in resting-state (black) and sleep-state (gray) acquisitions from the PSU dataset.

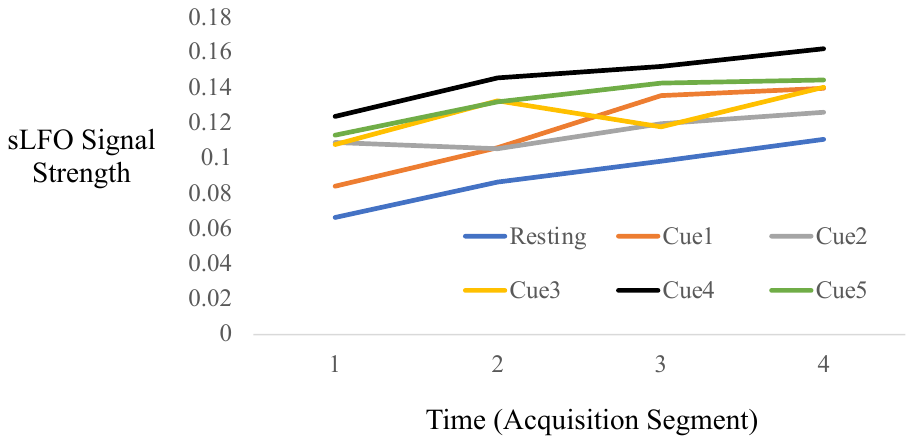
**a)**

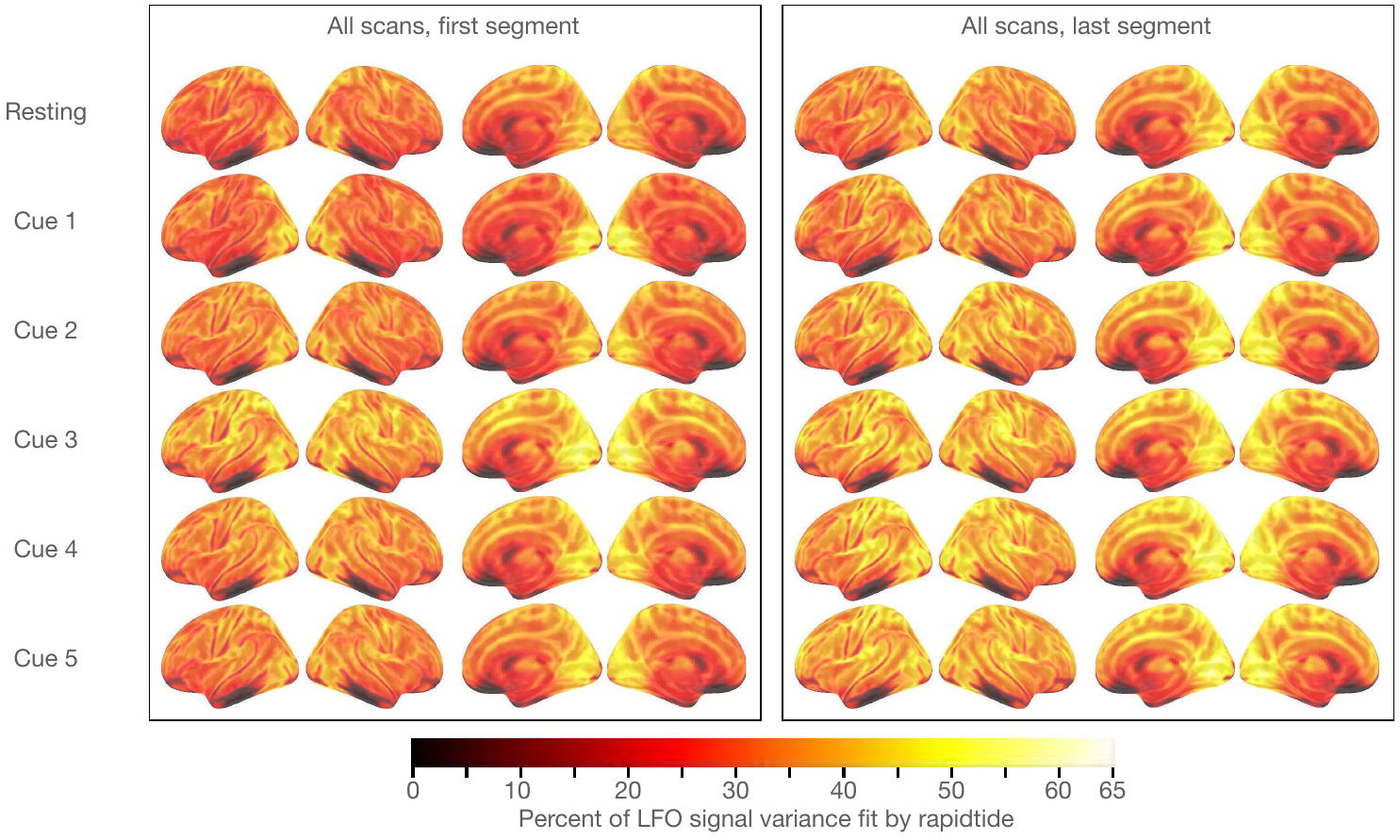

**b)**

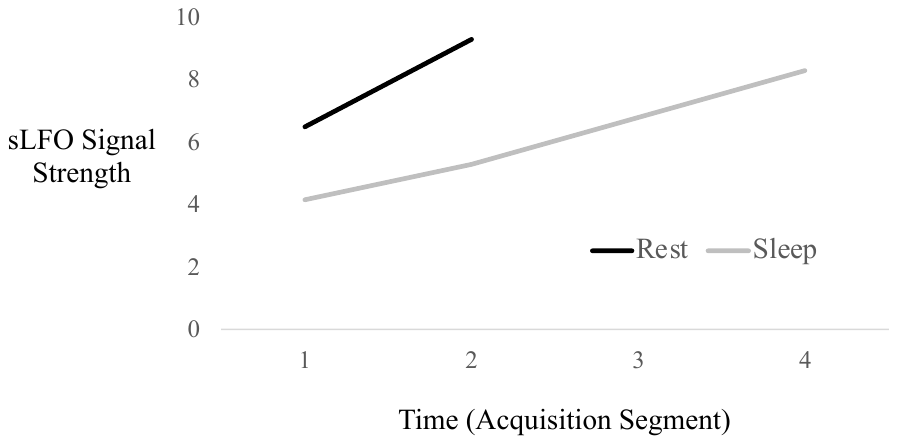

**c)**

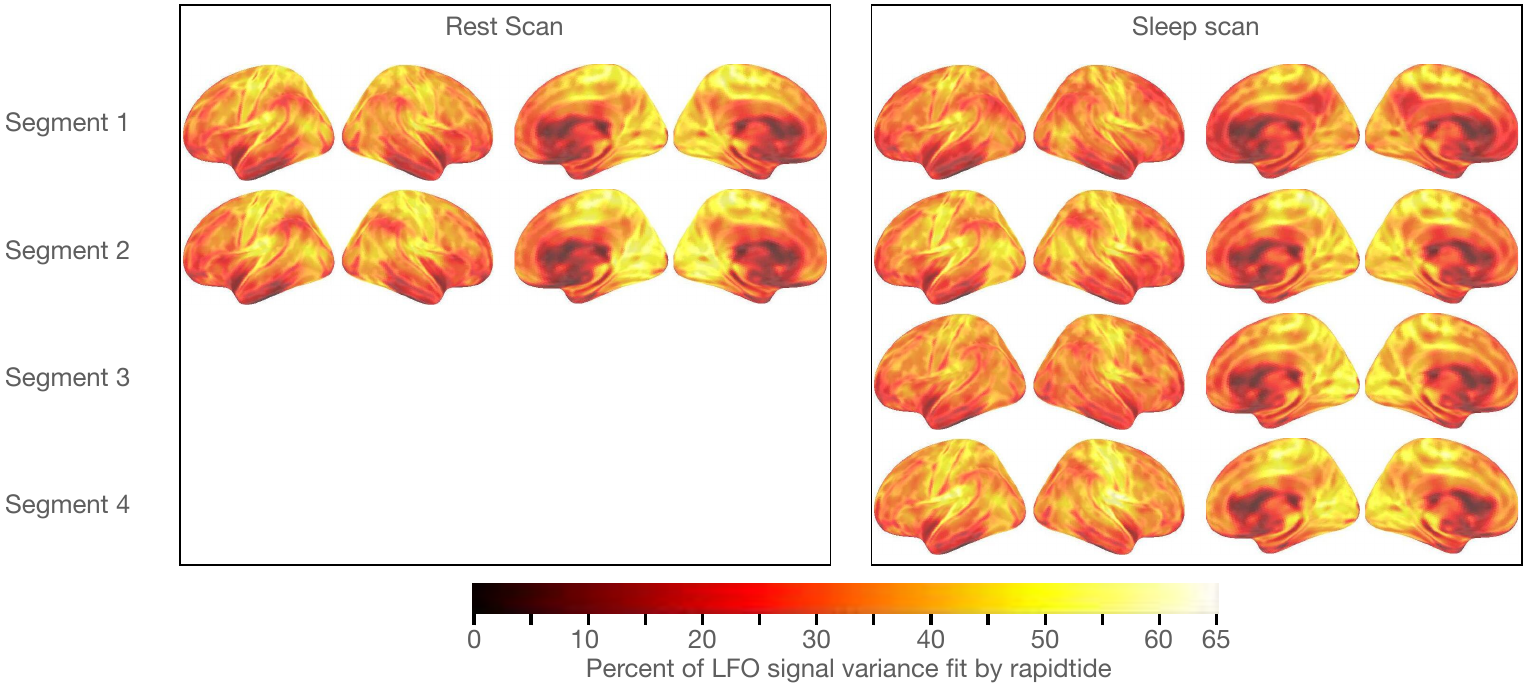

**d)**

**Figure S2.** Temporal and spatial properties of the sLFO signal in the MIC and PSU datasets. Brain-wide average percent of low frequency (0.01-0.15Hz) BOLD variance that was attributable to the non-neuronal sLFO signal at sequential time points in the a) MIC dataset and c) PSU dataset. Voxel-wise spatial distribution of the percent of low-frequency BOLD variance attributable to the sLFO signal during each temporal segment of the b) MIC dataset and d) PSU dataset.

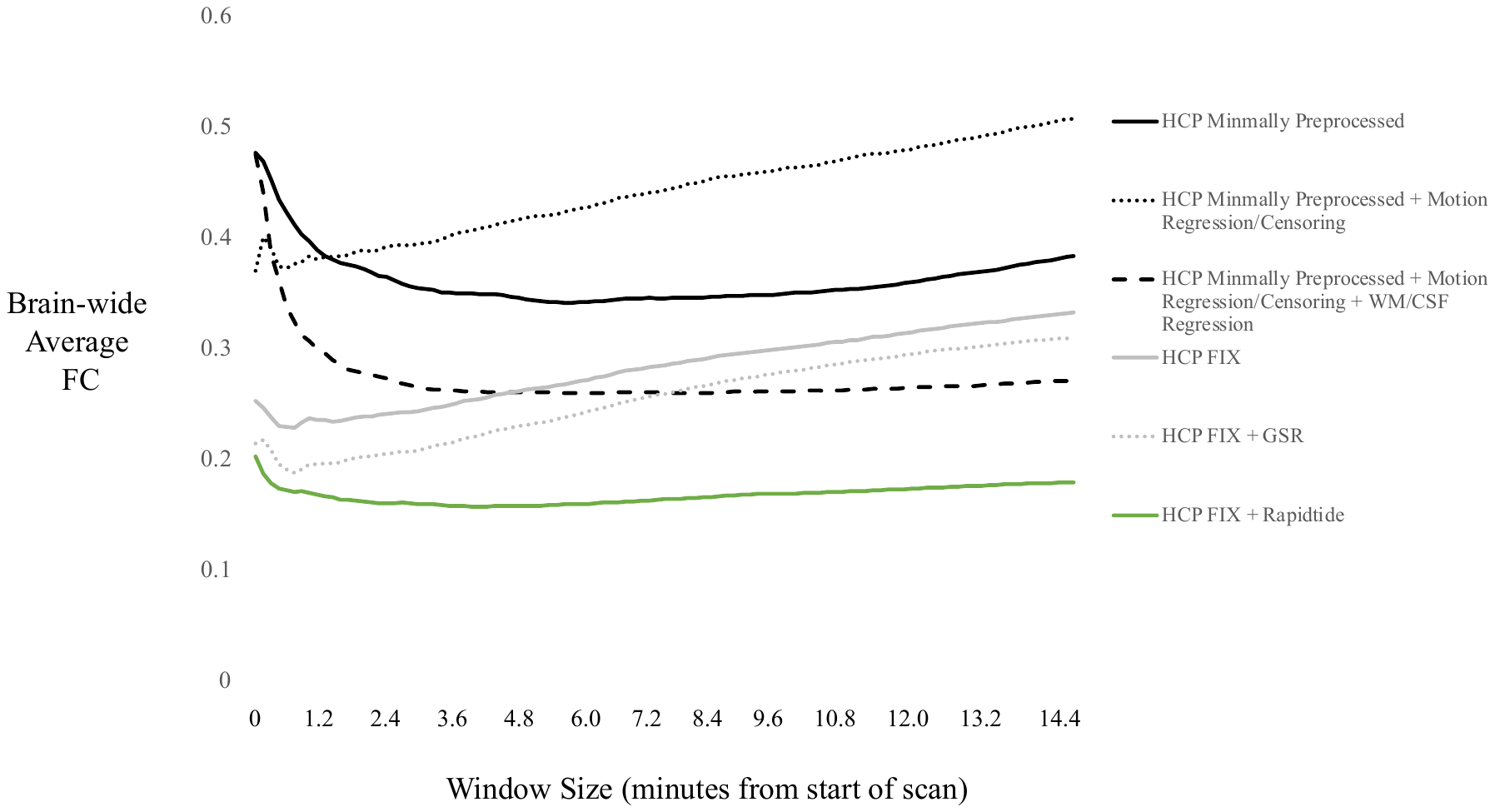

**Figure S3.** Group-averaged, mean brain-wide FC (*z*-score) computed over successively larger time windows for REST1 LR data with different preprocessing and denoising steps applied. Complementing FIX ICA with RIPTiDe denoising of the sLFO signal optimizes within-acquisition stability of brain-wide FC.

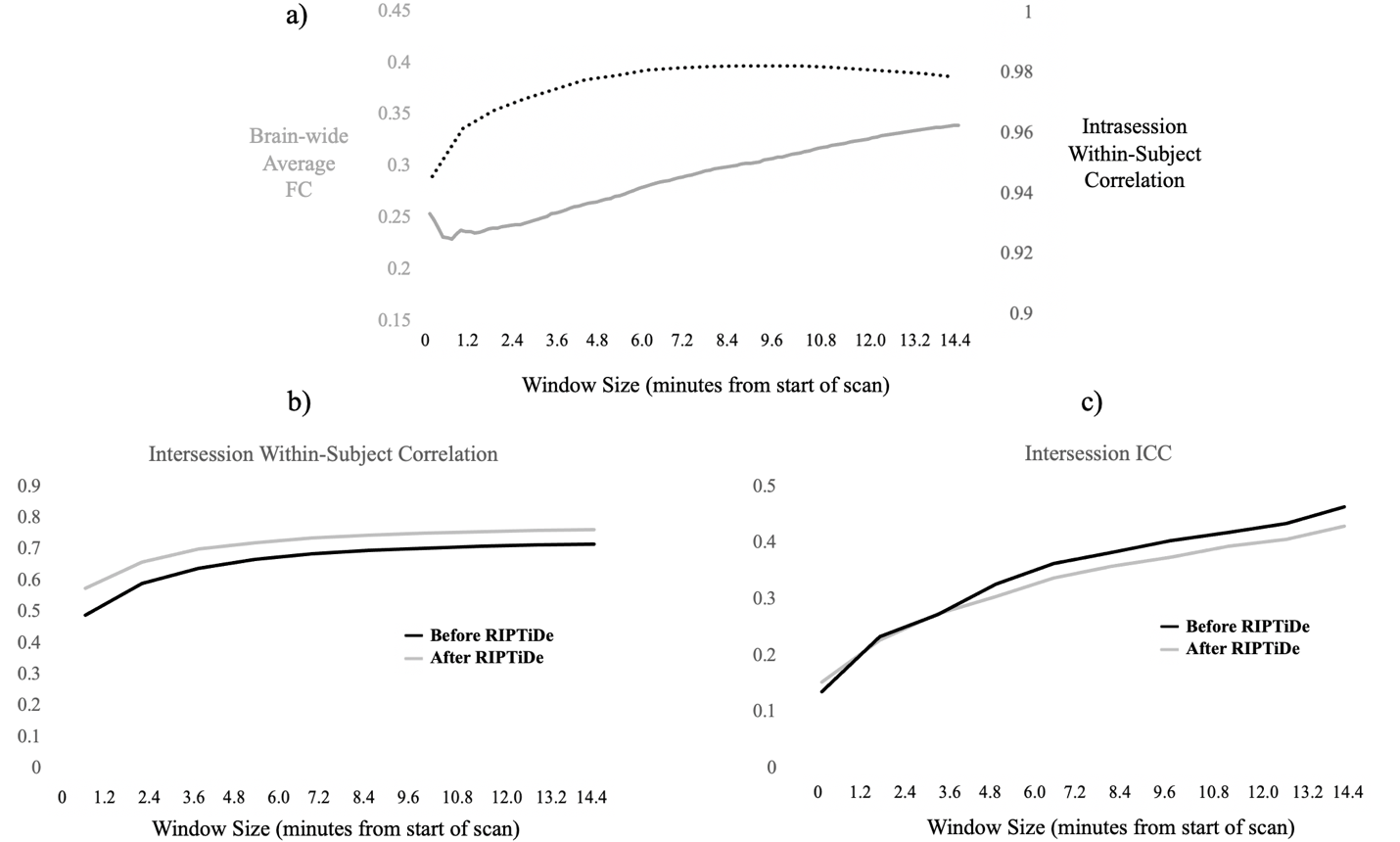

**Figure S4.** Within-scan reliability was largely independent of between-scan reliability and subject discriminability. a) Between-acquisition reproducibility (i.e., the group average correlation between a subject’s REST1 LR connectivity matrix and their REST1 RL connectivity matrix; dashed line) remained high (i.e., above 93%) and plateaued after only six minutes. Conversely, sLFO-driven FC inflation occurred throughout the scan (solid line). RIPTiDe denoising of the sLFO signal produced only small changes in b) between-scan reproducibility (i.e., the group average correlation between a subject’s REST1 LR connectivity matrix and their REST2 LR connectivity matrix) and c) ICC-indexed subject discriminability (ICC(2,1) using REST1 LR and REST2 LR).
